# The First Mitotic Division of the Human Embryo is Highly Error-prone

**DOI:** 10.1101/2020.07.17.208744

**Authors:** Emma Ford, Cerys E. Currie, Deborah M. Taylor, Muriel Erent, Adele L. Marston, Geraldine M. Hartshorne, Andrew D. McAinsh

**Author notes:** these authors contributed equally (Emma Ford, Cerys E. Currie).

## Abstract

Aneuploidy in human embryos is surprisingly prevalent and increases drastically with maternal age, resulting in miscarriages, infertility and birth defects. Frequent errors during the meiotic divisions cause this aneuploidy, while age-independent errors during the first cleavage divisions of the embryo also contribute. However, the underlying mechanisms are poorly understood, largely because these events have never been visualised in living human embryos. Here, using cell-permeable DNA dyes, we film chromosome segregation during the first and second mitotic cleavage divisions in human embryos from women undergoing assisted reproduction following ovarian stimulation. We show that the first mitotic division takes several hours to complete and is highly variable. Timings of key mitotic events were, however, largely consistent with clinical videos of embryos that gave rise to live births. Multipolar divisions and lagging chromosomes during anaphase were frequent with no maternal age association. In contrast, the second mitosis was shorter and underwent mostly bipolar divisions with no detectable lagging chromosomes. We propose that the first mitotic division in humans is a unique and highly error-prone event, which contributes to fetal aneuploidies.

## Introduction

Human reproduction requires the generation of eggs and sperm (haploid gametes) through two meiotic cell divisions. Fusion of gametes during fertilisation produces a diploid zygote (fertilised egg) that can then undergo the serial mitotic divisions required to build an embryo and ultimately a human being. The failure of homologous chromosomes or sister chromatids to segregate equally during female meiosis leads to deviations in chromosome number (Hassold and Hunt, 2001). Meiotic aneuploidies in human oocytes follow a U-shaped curve, with aneuploidy rates increasing exponentially with maternal age from about 35 years old (Gruhn et al., 2019). Studies of aneuploidy characteristics identify the precocious separation of sister chromatids and reverse segregation (two sister chromatids segregate) during meiosis I as errors which increase with maternal age. These errors are thought to be caused by a weakening in the cohesin complex, persistence of chromatid threads which hold sisters together and/or the geometry of kinetochore attachment (Gruhn et al., 2019; Patel et al., 2015; Zielinska et al., 2015). However, over 50% of blastocysts have blastomeres with different chromosome compositions (Chavez et al., 2012; McCoy, 2017). This “mosaicism” in early embryos is attributed to errors during the mitotic divisions, because errors in meiosis would affect all blastomeres. This mitotic-origin mosaicism is not necessarily lethal, and birth of mosaic babies following transfer of mosaic embryos is rare because affected cells can be selected against. However, some individuals can be born with placental, somatic or germ line mosaicism (Biesecker and Spinner, 2013; Kahraman et al., 2020). However, aneuploidies are clearly associated with infertility, miscarriages and developmental disorders (Lathi et al., 2008; Vitez et al., 2019). It is therefore crucial to establish the molecular origin of mitotic errors (as well as meiotic errors) and identify potential clinical risk factors and biomarkers that could stratify embryos for transfer to patients. Non-invasive imaging of human embryos has shown that the timing of early mitotic divisions can, to some extent, predict whether or not an embryo will develop to the blastocyst stage (Wong et al., 2010), and correlates with aneuploidy (Chavez et al., 2012). However, a limitation of these studies is that they do not directly follow chromosome segregation and assign error-rates to each mitotic division. Here we have established live-embryo “chromosome imaging” throughout the first two mitotic cleavage divisions of the human embryo using SiR-DNA, a far-red fluorogenic probe for DNA (Lukinavicius et al., 2015). Our experiments provide initial insights into the timing and fidelity of chromosome segregation during the human embryonic divisions that mark the switch from meiotic to mitotic cell division programmes.

## Results & Discussion

Human embryos for this study were provided by women undergoing intracytoplasmic sperm injection (ICSI) or in vitro fertilisation (IVF). We received embryos that had been deselected from further use in patient treatment because they contained either more or fewer than the normal 2 pronuclei (PN; as expected for a correctly fertilised oocyte), or were still at the metaphase II (MII) stage and appeared to be unfertilised when clinical decisions were taken). By the start of imaging experiments 64% of all embryos where NEB was visualised contained 2PN, indicative of delayed pronuclear formation. For live-embryo imaging, the glycoprotein-rich zona pellucida was removed before addition of SiR-DNA, thus allowing visualisation of chromosomes (far red) and the whole embryo (bright field). By using long-term time-lapse microscopy (every 15 or 10 mins for 24-36 hours) and acquiring a 90 µm z-stack in both channels, we could determine the timing of key cell division and chromosome segregation events (Fig. 1a,b).

**Figure 1:**
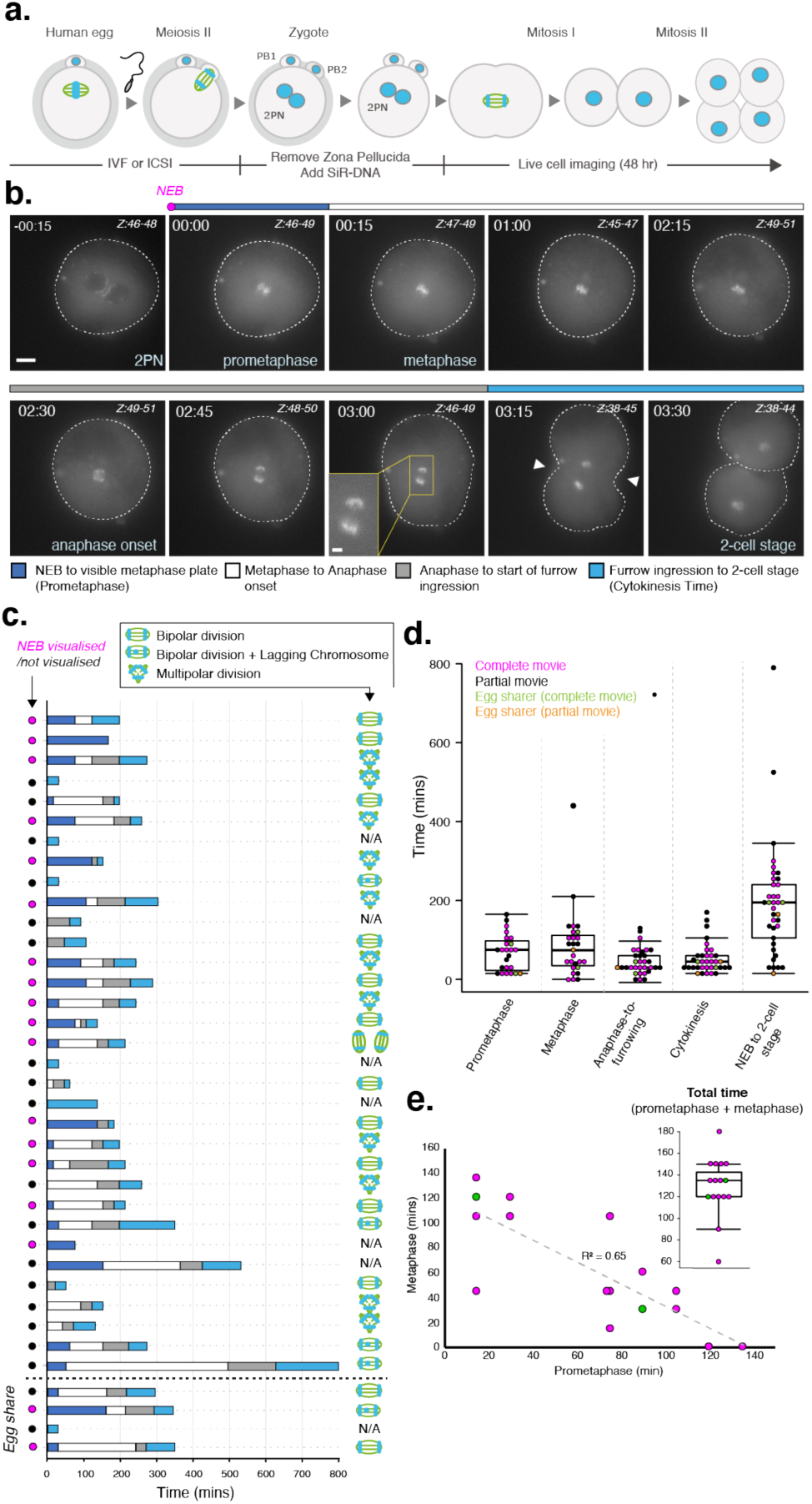
Live cell imaging of chromosome dynamics during mitosis I of human embryos. (a) Schematic outlining key steps in embryo preparation for chromosome imaging. (b) Time lapse imaging of a representative human embryo progressing through mitosis I. Chromosomes were visualised using SiR-DNA dye. Z indicates slices shown as maximum intensity projection. Time in hours:mins, scale bar 20*μm.* White arrows indicate onset of cleavage furrow ingression. (c) History plots of all human embryos imaged during this study. Coloured bars denote timings of critical stages during mitosis I and cartoons indicate phenotypes observed. Pink/black dots indicate whether NEB was visualised during filming. (d) Quantification of data presented in (c) to compare timings of each phase during mitosis I. (e) Correlation of metaphase vs prometaphase in embryos where the stages were filmed in their entirety, R^2^ = 0.65. Inset box and whisker plot shows the total duration of both stages for each embryo.

Using our live-embryo imaging movies we measured the time from nuclear envelope breakdown (NEB) of pronuclei to the appearance of a metaphase plate (prometaphase time) and the time from metaphase plate formation to the onset of sister chromatid separation (metaphase time) (Fig. 1c). The time taken to complete these events reflects the efficiency of key biological events. During prometaphase, the microtubule-based mitotic spindle assembles and chromosomes are captured, bi-oriented and aligned on the metaphase plate. The subsequent metaphase time reflects maturation of kinetochore-microtubule attachments and release of spindle checkpoint mediated inhibition of the anaphase promoting complex/cyclosome (APC/C), thus allowing anaphase onset. The median times for prometaphase and metaphase were 75 mins (n=17) and 75 mins (n=24), respectively (Table S1, Fig. 1d). NEB was not visualised in 19 embryos (Fig. 1c) (hence the deviation in n) and their prometaphase time is potentially underestimated, though values nevertheless fall within the distribution (Fig 1d, black *vs* pink dots). The interquartile range (IQR) indicates that transition through these two phases is highly heterogeneous (67.5 and 77.5, respectively). Interestingly, plotting metaphase time versus prometaphase time for each embryo revealed an inverse relationship (Fig. 1e), suggesting that a timing mechanism operates to fix the time from NEB to anaphase onset at ∼2.2 hours. This has implications for the functioning of surveillance mechanisms (see below). We next measured the time from anaphase onset to the first indication of furrow ingression, and from that point to the completion of cell division (2-cell stage). The median times were 30 mins (n=31; IQR = 30) and 45 mins (n=35; IQR = 30), respectively (Fig 1d). Again, these phases are much longer than events measured in human somatic cells (∼7 mins from anaphase onset to 2-cell stage (Spira et al., 2017); although not as variable as pre-anaphase events. Taking all phases together from NEB to 2-cell stage, the total duration of the first embryonic mitosis was 195 mins (n=37; IQR = 135) (Fig 1d). This is consistent with observations in both *Xenopus* egg extracts and mouse embryos showing that the first embryonic mitosis is extended (Ajduk et al., 2017; Chesnel et al., 2005; Sikora-Polaczek et al., 2006).

The extended duration of prometaphase and metaphase in the first human embryonic mitosis suggests that the process of spindle assembly and formation of kinetochore interactions are intrinsically inefficient. Indeed, 38% of embryos (n=12) failed to form a bipolar spindle at all, and underwent a multipolar division separating DNA into three or more masses (Fig. 2a). This multipolar chromosome segregation was also poorly coordinated with cytokinesis as resultant DNA masses were not always associated with a daughter cell (Table S2). We note that mis-fertilisation (3PN embryos) does not provide the only explanation for multipolar divisions as some 2PN embryos also divided in a multipolar fashion. Embryos with multipolar spindles – and therefore an increased frequency of mal-oriented kinetochore-microtubule attachments – did not delay before anaphase compared to those with bipolar spindles (Fig. 2b). This implies that the spindle checkpoint (which normally functions to delay anaphase onset until all kinetochore-microtubule attachments are corrected) is weakened, non-functional, or has a finite lifetime, after which cells bypass the checkpoint without correcting kinetochore-microtubule attachment errors.

**Figure 2:**
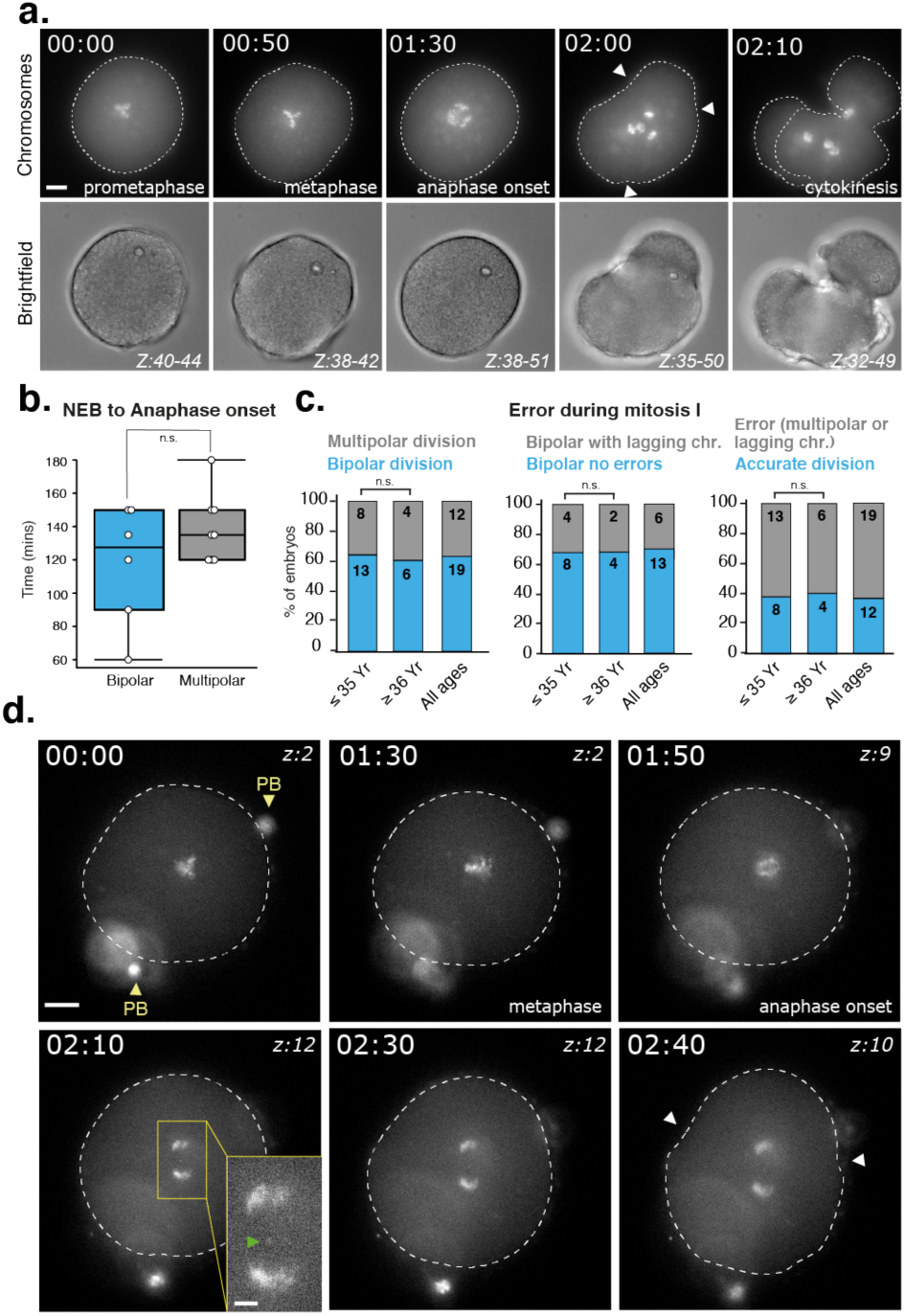
The first mitotic division of human embryos is heterogeneous. (a) Time lapse imaging of a human embryo undergoing a multipolar mitosis I resulting in multiple daughter cells. Z indicates slices shown as maximum intensity projection. Chromosomes were visualised using SiR-DNA dye. White arrows indicate onset of cleavage furrow ingression. Time in hours:mins, scale bar 20μm. (b) Quantification of the duration of NEB to anaphase onset in bipolar vs multipolar divisions. P value from a Mann Whitney U test. (c) Embryos were analysed for the presence of mitotic errors (lagging chromosomes and multipolar divisions). N numbers are presented within each bar. Significance calculated using a Fisher’s exact test. (d) Same as (a) except this embryo undergoes mitosis I in the presence of a lagging chromosome (green arrow), single Z slices displayed. PB: Polar bodies. scale bar 20μm, inset 5μm.

Following anaphase onset in embryos that underwent a bipolar division, we also observed a high incidence (31%) of lagging chromosomes (Fig. 2c,d). Studies in human somatic cells have shown that such events are indicative of merotelic kinetochore-microtubule attachments that failed to correct before anaphase onset (Cimini et al., 2002). These attachments are not monitored by the spindle checkpoint and are a likely source of nondisjunction events (Gregan et al., 2011). The fraction of embryos that exhibited a lagging chromosome or a multipolar division was 61% (Fig. 2c). Because errors from a mitotic origin are reported to be independent of maternal age (McCoy, 2017), we next tested whether the frequency of lagging chromosomes or multipolar spindles at the first mitotic division were associated with older patients (>36 years) *vs.* younger patients (<35 years), as 35 years seems to be the age at which meiotic errors increase exponentially (Gruhn et al., 2019). The incidence of either multipolar spindles or lagging chromosomes appeared to be age-independent (Fig. 2c). Taken together, our live cell imaging of chromosomes reveals how the first mitotic division of the human zygote is inefficient and highly error-prone.

One potential limitation of these observations is the use of embryos that were delayed or unsuitable for further treatment of patients (deselected embryos). The gold standard would be normal 2PN human embryos. To access such embryos, we established an egg sharing programme whereby patients aged up to 32 years, undergoing IVF or ICSI and anticipated to produce plentiful eggs in response to stimulation, voluntarily elect to share half of those eggs with the research programme. Such patients receive their treatment for a reduced cost subsidised by the research funder. This is in keeping with UK law and approved by the NHS Research Ethics Committee and the Human Fertilisation and Embryology Authority. Ten oocytes were recovered from a patient aged 29 with infertility caused by a combination of blocked Fallopian tubes and a male factor. The patient had achieved a previous successful pregnancy through ICSI. The oocytes designated to research were fertilised (by ICSI using donor sperm) giving rise to five embryos (all 2PN) that were then imaged for ∼2 days. Four of these embryos (one did not survive removal of the zona pellucida) underwent anaphase onset with a bipolar spindle (Fig. 3a), although in two embryos lagging chromosomes were detected at anaphase (Fig. 3b). Moreover, the timing of key mitotic events in these shared embryos fell within the distributions of the deselected research embryos (Fig. 1c and green/orange data points in Fig. 1d,e). Therefore, the presence of errors and extended mitotic and cytokinesis durations are true characteristics of mitosis I and not simply a consequence of using clinically deselected embryos for research.

**Figure 3:**
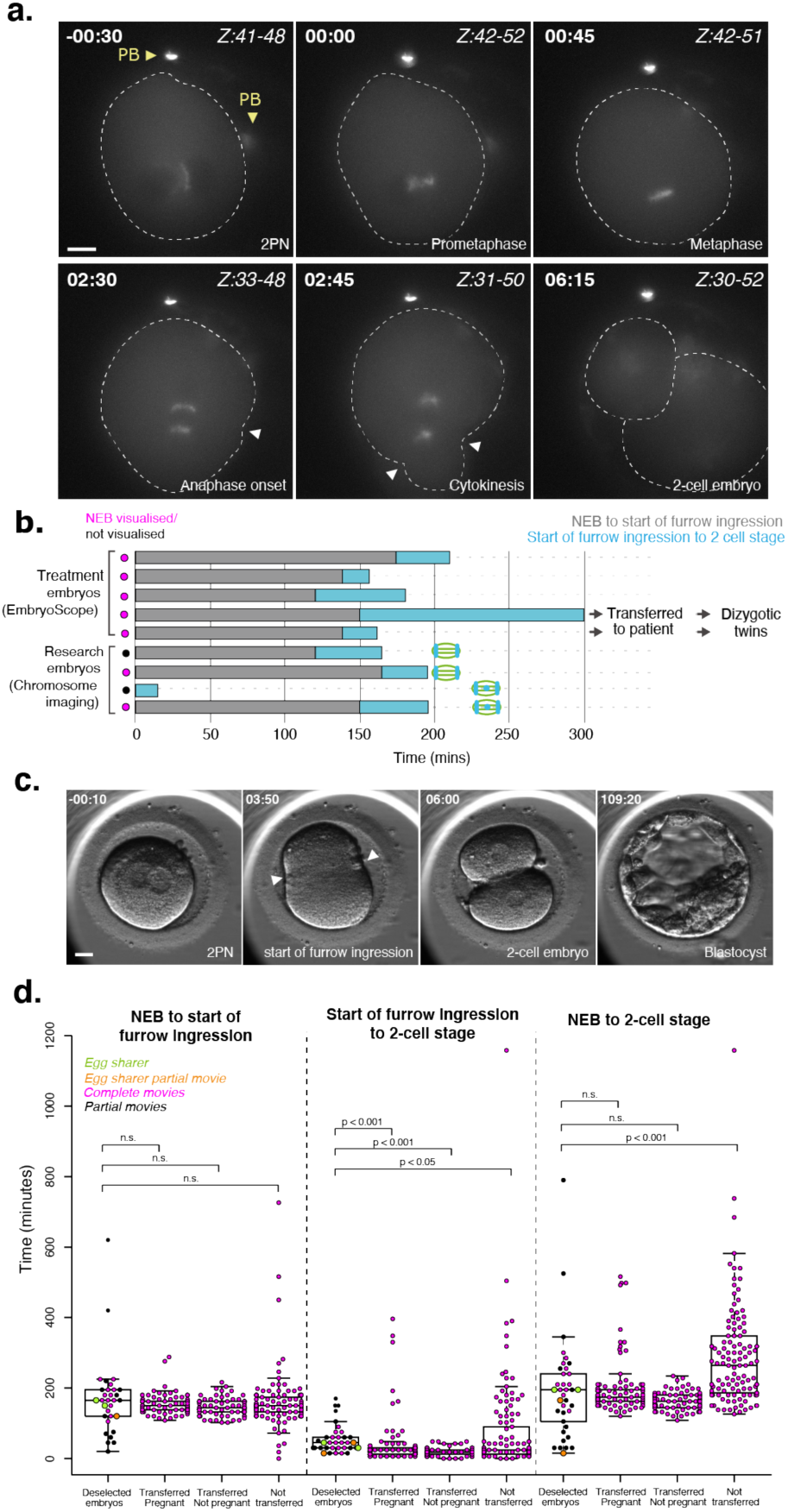
Freshly-shared research embryos are consistent with deselected embryos. (a) Time lapse imaging showing a representative human embryo donated by an egg-share-to-research patient undergoing mitosis I. PB:Polar Bodies, Z indicates which slices are shown as maximum intensity projection. White arrows denote the onset of cleavage furrow ingression. Chromosomes were visualised using SiR-DNA dye. Time in hours:mins, scale bar: 20μm. (b) History plots of egg share embryos; half used for patient treatment and half donated to research. Coloured bars denote timings of critical stages during mitosis I, visualised by brightfield illumination on an EmbryoScope™ or widefield microscope. Pink/black dots indicate whether NEB was visualised during filming. (c) Representative movie stills showing EmbryoScope™ monitoring of an egg-share-to-research embryo used for patient treatment. White arrows denote onset of cleavage furrow ingression. Scale bar: 20μm. (d) Quantification of timings of key events during mitosis I, of both research embryos imaged on a widefield microscope (with SiR-DNA staining) and indicated groups of treatment embryos monitored using the EmbryoScope™. P-values from a Mann-Whitney U test.

Because half the sharer’s eggs were imaged in the clinic by time lapse microscopy to inform the selection of embryos for transfer to the patient, we were able to compare autologous timings directly with our live-cell imaging. The Hoffmann contrast images from the clinic allow determination of the timing of pronuclear fading (equivalent to NEB), initiation of furrow ingression and formation of daughter cells (chromosomes are not visible by this method) (Fig. 3c). Fig. 3b shows that the durations between NEB, furrow ingression and 2-cell stage were similar to the research embryos. Furthermore, two of the sharer’s embryos were transferred resulting in a dizygotic twin clinical pregnancy, confirmed by fetal heart activity on ultrasound scan at 7 weeks of gestation. This again shows how the cell division timings, and potentially the presence of lagging chromosomes reported here in deselected embryos, can be compatible with embryo development and implantation.

The main limitation of the egg sharer data is that it is limited to one patient. We therefore sought to compare our pooled chromosome imaging data (including both shared and deselected embryos) to the timing of events in a cohort of human embryos transferred singly that gave rise to live birth. In addition, we analysed deselected (not transferred or cryopreserved) embryos from the same treatment cycles where successful pregnancies were achieved, including those which underwent apparently normal first and second cleavage divisions or those which underwent an abnormal first cleavage division. As a further control, we also analysed movies of embryos that were transferred singly into a further group of patients who did not become pregnant. The timing from NEB to the start of furrow ingression in the deselected research embryos (which underwent chromosome imaging) vs. transferred pregnant, non-pregnant or non-transferred embryos were broadly similar (Fig. 3d). These results show that data collected from chromosome imaging of research embryos are largely consistent with embryos that generate normal pregnancies. These data also show how SiR-DNA treatment does not perturb embryo progression. The median mitosis I timings were comparable between the transferred pregnant population and our research embryos (174 vs 195 mins). We also note how long mitosis I times can be compatible with pregnancy. Thus, we consider that our data are broadly representative of the normal situation, despite the material being deselected for clinical use.

Following the first mitotic division we continued to film embryos to capture the second mitosis (Fig. 4a, Fig. S1). We observed 19 cases, including the egg sharer embryos, and determined the timing of key cell division events as described above for the first mitotic division (Fig. 4b; Table S3). We found that the NEB to 2 cell duration of mitosis II was significantly shorter than that of mitosis I (210 mins (IQR = 48.8) vs 150 (IQR = 45) p = 0.004) (Fig. 4c). This reduction in duration of mitosis II was mainly due to a decrease in cytokinesis time (15 mins, IQR=5 min *vs* 45 min IQR=30 min; p = 5.3 ×10^−4^). For embryos imaged undergoing both mitosis I and mitosis II, the elapsed time between the 2-cell stage and 4-cell stage was 15.5 hr (n=9, IQR = 1.8), which is very similar to the timings reported in embryos used for patient treatment (Cruz et al., 2012) (Figure S1). Importantly, we observed zero lagging chromosomes during anaphase of mitosis II, and only two multipolar divisions (10%). This further demonstrates that our findings for the first division were not a consequence of the imaging and culturing conditions. Moreover, it suggests that the first mitotic division is the major contributor to mitotic aneuploidies in humans.

**Figure 4:**
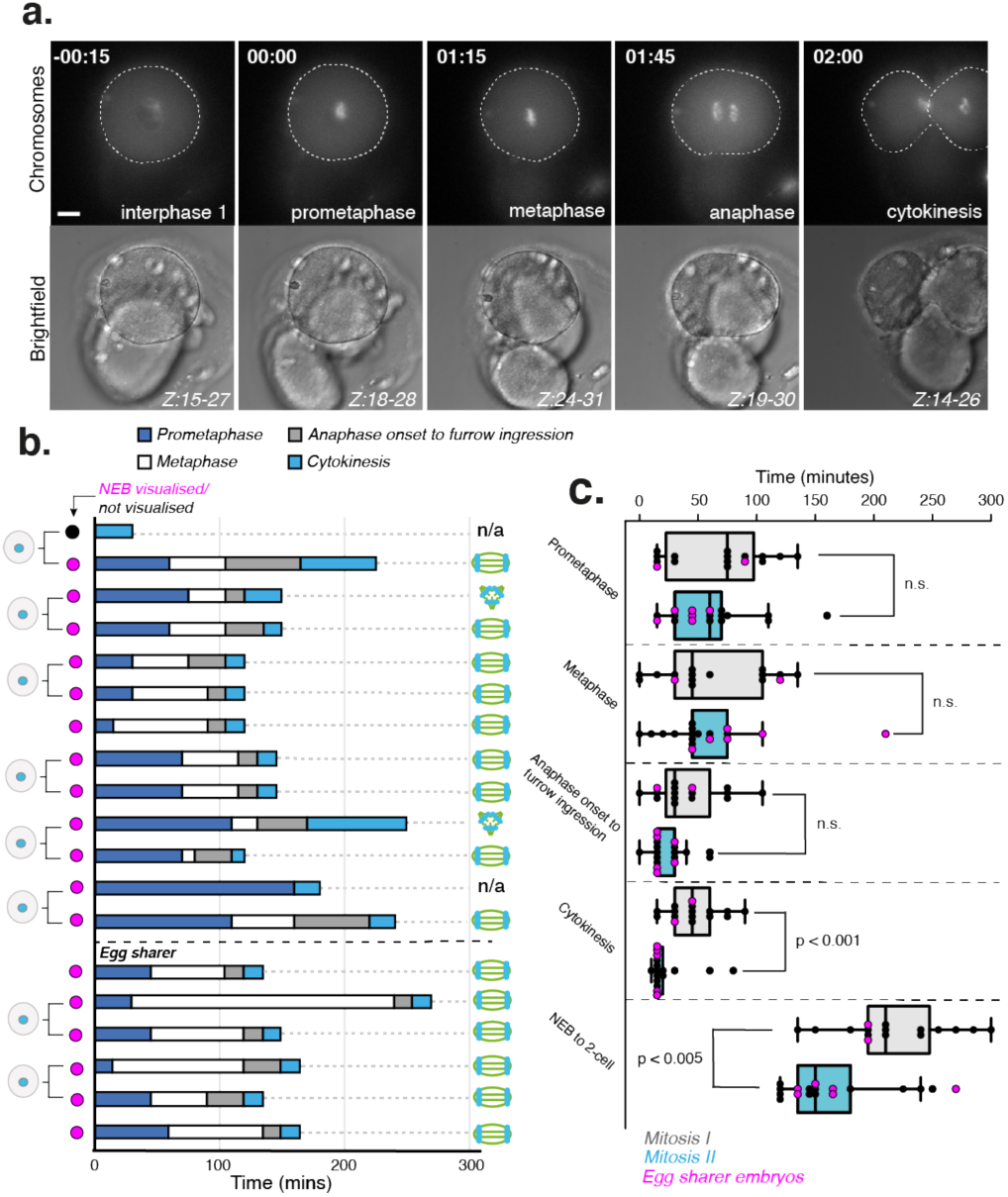
The second mitotic division of human embryos is shorter and less error-prone. (a) Time lapse imaging of a representative human embryo progressing through mitosis II. Chromosomes were visualised using SiR-DNA dye. Z indicates slices shown as maximum intensity projection. Time in hours:mins, scale bar: 20μm. (b) History plots of all human embryos undergoing mitosis II imaged for this study. Coloured bars denote timings of critical stages during mitosis II and spindle cartoons indicate phenotypes observed. Daughter cell cartoons indicate cells arising from the same mother embryo. (c) Quantification of key events during mitosis I (n=16) and II (n=18), of research embryos where the process was filmed in its entirety. P values from a Mann-Whitney U test.

What is the origin of the errors we observe during the first mitotic division? Work in murine and bovine embryos has shown that a distinct paternal and maternal spindle form and then fuse together during the first cleavage division (Destouni et al., 2016; Reichmann et al., 2018). This distinctive spindle formation distinguishes mitosis I from the later mitoses and could explain why it is so error-prone (the two spindles not fusing together properly could cause multipolar spindles and an increased prevalence of merotelic attachments). A dual spindle has also been reported in a human zygote (Xu et al., 2019). We were able to observe the formation of a dual spindle in a 3PN embryo (Fig. S2), providing further evidence that human embryos can assemble such a structure during the first embryonic mitosis. Our data also provide evidence for a timer mechanism that sets a prometaphase-metaphase duration to allow sufficient time for spindle assembly and chromosome bi-orientation. This may be necessary if, like in mouse, the spindle checkpoint is dispensable during the initial embryonic divisions (Dobles et al., 2000). Moreover, the spindle checkpoint proteins are displaced from kinetochores well before formation of the metaphase plate during the first, but not the second, mitosis in mouse embryos (Sikora-Polaczek et al., 2006). This is in line with our observations that the first, but not second, division is error-prone. However, at this stage we cannot rule out that kinetochore-microtubule attachments are monitored to some extent. The nature of the timing mechanism that regulates the duration of mitosis I is unknown, although experiments in mouse suggest that delayed activation of APC/C by polo kinase 1 prolongs NEB to anaphase onset independently of kinetochore-microtubule attachment status (Ajduk et al., 2017). Alternatively, the mechanism may be analogous to the way somatic cells set a minimal pre-anaphase time using checkpoint protein complexes (Meraldi et al., 2004; Rodriguez-Bravo et al., 2014).

Finally, this study documents the first comparison of chromosome segregation during the first and second cleavage divisions in routinely deselected human research embryos vs. freshly shared human embryos. The similarities between the two provide firm ground for the use of deselected material in the study of human developmental mechanisms, and reveal how the first mitotic division is the major source of mitotic-origin aneuploidies in human embryos.

## METHODS

### Human Embryos

For donation of human embryos to research: The NHS Research Ethics Committee approved both the research project (04/Q2802/26) and egg sharing to research (19/WM/0003). All work was conducted under a Research Licence from the Human Fertilisation and Embryology Authority (HFEA; R0155). Informed consent for donation of eggs, embryos and sperm to research was provided by patients undergoing in vitro fertilisation (IVF) or intracytoplasmic sperm injection (ICSI) at the Centre for Reproductive Medicine (CRM), University Hospitals Coventry and Warwickshire (UHCW) NHS Trust. The material used for research was unsuitable for the patient’s treatment and would otherwise have been disposed or was collected from a volunteer woman undergoing egg sharing to research. Mature (metaphase II) eggs collected from the egg sharer were inseminated with fertile donor sperm using ICSI during the course of the research.

### Live Cell Imaging

Research oocytes and embryos were collected from the clinic around 5 hours after clinical decision making. During this time, some embryos had progressed to later stages. The zona pellucida of individual embryos was removed by brief incubation and pipetting in prewarmed acid Tyrode’s solution (Sigma). Embryos were transferred to a Fluorodish (WPI) containing 2 µM SiR-DNA (Spirochrome) diluted in prewarmed Cleav media (Origio) under mineral oil. Embryos were transported about 14 km from UHCW CRM to Warwick Medical School (WMS) in a portable incubator (K Systems) held at 37°C, and imaged immediately upon arrival. Image stacks (60 x 1.5 μm optical sections; 1×1 binning) were acquired every 10 or 15 minutes for a 24-36 hr period with a 40x oil-immersion 1.3 NA objective using an Olympus DeltaVision Elite microscope (Applied Precision, LLC) equipped with a CoolSNAP HQ2 camera (Roper Scientific). Images were acquired at 32% neutral density using Cy5 filter and an exposure time of 0.05s. A stage-top incubator (Tokai Hit) maintained embryos at 37°C and 5% CO_2_ with further stabilisation from a microscope enclosure (Weather station; Precision Control) held at 37°C. The temperature was confirmed with a calibrated probe (Fluke 52). Image sequences were inspected and analysed by hand using OMERO (Open Microscopy Environment).

### Clinical Imaging of Human Embryos

298 human embryos were imaged on an EmbryoScope™ for patients who received ICSI treatment. Images were collected every 10 mins for up to 6 days and the timing of events (NEB, the start of cytokinetic furrow ingression and the appearance of 2 distinct cells) during the first cell division determined. These embryos fall into 3 categories: transferred embryos which gave rise to pregnancy and live birth, transferred embryos which did not give rise to pregnancy and embryos from the same patients who did become pregnant, but which were not transferred due to abnormal morphology at embryological examination.

### Data and Statistical Analysis

Mann-Whitney U tests for Fig. 2b, 3d and 4c and Fisher’s exact tests for Fig. 2c were performed using MATLAB R2020A (Mathworks) inbuilt functions. Box-and-whisker plots show the IQR and minimum/maximum values, plus all individual data points.

### CRediT

**Emma Ford:** Investigation, Data curation, Formal Analysis, Editing, Methodology, **Cerys Currie**: Investigation, Data curation, Formal Analysis, Reviewing and Editing, Validation, **Muriel Erent**: Data curation, Visualisation, **Debbie Taylor:** Methodology, Resources, **Adele Marston**: Conceptualization, Reviewing and Editing, **Geraldine Hartshorne:** Conceptualization, Writing, Reviewing and Editing, Supervision, Ethical approvals, **Andrew McAinsh** Conceptualization, Supervision, Formal Analysis, Original draft preparation, Writing and Editing.

## ACKNOWLEDGEMENTS

We are hugely thankful and indebted to the patients, embryologists and research nurses at the Centre for Reproductive Medicine, University Hospitals Coventry and Warwickshire NHS Trust. We also gratefully acknowledge CAMDU (Computing and Advanced Microscopy Unit) for their support and assistance in this work. A.D.M, A.L.M., G.M.H., C.C. and D.T. are supported by a Wellcome Collaborative Award (215625). E.F. is supported by a Warwick Collaborative Postgraduate Research Scholarship with UHCW. A.D.M is also supported by a Wellcome Senior Investigator Award (106151) and a Wolfson Royal Society Research Merit Award (WM150020). G.M.H. is also supported by the WPH Charitable Trust. A.L.M. is also supported by a Wellcome Senior Research Fellowship (107827) and core funding for the Wellcome Centre for Cell Biology (203149).

---

**Table S1:** Mitosis I timings.

| Patient number | NEB? | Prometaphase | Metaphase | Anaphase onset to furrow ingression | Cytokinesis | NEB to 2-cell |
| --- | --- | --- | --- | --- | --- | --- |
| 3000 | No | 50 | 440 | 130 | 170 | 790 |
| 3004 | No | 60 | 90 | 70 | 50 | 270 |
| 3028 | No | n/a | 40 | 30 | 60 | 130 |
| 3034 | No | n/a | 90 | 30 | 30 | 150 |
| 3045 | No | n/a | n/a | 20 | 30 | 50 |
| 3067 | No | 150 | 210 | 60 | 105 | 525 |
| 3068 (i) | YES | 75 | n/a | n/a | n/a | 75 |
| 3068 (ii) | No | 30 | 90 | 75 | 150 | 345 |
| 3083 * | YES | 15 | 135 | 30 | 30 | 210 |
| 3101 | No | n/a | 135 | 60 | 60 | 255 |
| 3148 * | YES | 15 | 45 | 105 | 45 | 210 |
| 3158 * | YES | 15 | 105 | 30 | 45 | 195 |
| 3159 (i) * | YES | 135 | 0 | 30 | 15 | 180 |
| 3159 (ii) | No | n/a | 0 | 0 | 135 | 135 |
| 3169 | No | n/a | 15 | 30 | 15 | 60 |
| 3170 | No | n/a | n/a | n/a | 30 | 30 |
| 3172 * | YES | 30 | 105 | 30 | 45 | 210 |
| 3181 * | YES | 75 | 15 | 15 | 30 | 135 |
| 3197 (i) * | YES | 30 | 120 | 45 | 45 | 240 |
| 3197 (ii) * | YES | 105 | 45 | 75 | 60 | 285 |
| 3201 * | YES | 90 | 60 | 30 | 60 | 240 |
| 3204 | No | n/a | n/a | 45 | 60 | 105 |
| 3209 | No | n/a | n/a | 60 | 30 | 90 |
| 3210 * | YES | 105 | 30 | 75 | 90 | 300 |
| 3215 | No | n/a | n/a | 0 | 30 | 30 |
| 3218 * | YES | 120 | 0 | 15 | 15 | 150 |
| 3226 (i) | No | n/a | n/a | n/a | 30 | 30 |
| 3226 (ii) * | YES | 75 | 105 | 45 | 30 | 255 |
| 3227 | No | 15 | 135 | 30 | 15 | 195 |
| 3233 (i) | No | n/a | n/a | n/a | 30 | 30 |
| 3233 (ii) * | YES | 75 | 45 | 75 | 75 | 270 |
| 3236 | YES | 165 | n/a | n/a | n/a | 165 |
| 3246 * | YES | 75 | 45 | 0 | 75 | 195 |
| 3247 (i) - egg share * | YES | 15 | 120 | 15 | 45 | 195 |
| 3247 (ii) - egg share | No | n/a | n/a | n/a | 15 | 15 |
| 3247 (iii) - egg share * | YES | 90 | 30 | 45 | 30 | 195 |
| 3247 (iv) - egg share | No | 15 | 75 | 30 | 45 | 165 |
| **Median** |  | **75** | **75** | **30** | **45** | **195** |
| **IQR** |  | **67.5** | **77.5** | **30** | **30** | **135** |

<sup>1</sup> \* denotes 16 embryos where mitosis I was filmed in its entirety

**Table S2:** Mitosis I phenotypes.

| Patient number | Bipolar/<br>multipolar | Lagging<br>chromosomes? | Maternal<br>age | Treatment | Number of chromosome masses formed | Number of cells formed |
| --- | --- | --- | --- | --- | --- | --- |
| 3000 | Bipolar | YES | 43 | IVF | 2 | 2 |
| 3004 | Bipolar | YES | 37 | ICSI | 3 | 2 |
| 3028 | Multipolar | NO | 35 | ICSI | 2 | 4 |
| 3034 | Multipolar | YES | 28 | IVF | 4 | 2 |
| 3045 | Bipolar | NO | 31 | IVF | 2 | 2 |
| 3067 | NA | YES | 23 | ICSI | DNA moved to the top of the cell, not visible | 2 |
| 3068 (i) | NA | n/a | 29 | ICSI | embryo moved out of the field of imaging | embryo moved out of the field of imaging |
| 3068 (ii) | Bipolar but one mass<br>stays on top | YES | 29 | ICSI | 3 | 2 |
| 3083 * | Bipolar | NO | 45 | IVF | embryo moved out of the field of imaging | 2 |
| 3101 | Multipolar | NO | 38 | IVF | 6 | 2 |
| 3148 * | Bipolar | NO | 38 | ICSI | 2 | 2 |
| 3158 * | Multipolar | NO | 30 | IVF | First 3 DNA which then fragment into at least 5<br>masses: 1 in one cell, 4 in the other | 2 |
| 3159 (i) * | Bipolar | NO | 38 | IVF | not clear, one big one that stays in one cell. | 2 |
| 3159 (ii) | NA | no anaphase<br>onset | 38 | IVF | remains in 2 masses the whole time, | 3 |
| 3169 | Bipolar | NO | 35 | IVF | 2 | 2 |
| 3170 | NA | n/a | 36 | IVF | SiR DNA did not work | 2 |
| 3172 * | Multipolar | YES | 39 | IVF | 5 | 3 |
| 3181 * | Bipolar | NO | 31 | IVF | 3 | 2 |
| 3197 (i) * | Multipolar | NO | 30 | IVF | 3 | many |
| 3197 (ii) * | Bipolar | NO | 30 | IVF | 3 main groups, but some smaller masses around | 3 |
| 3201 * | Multipolar | NO | 38 | IVF | 4, 1 in one cell, 3 in the other | 2 |
| 3204 | Bipolar | NO | 30 | IVF | 2 | 2 |
| 3209 | NA | n/a | 30 | IVF | DNA staining did not work properly | 2 |
| 3210 * | Multipolar | YES | 37 | IVF | >5 | 2 |
| 3215 | Bipolar | YES | 31 | IVF | 2 | 2 |
| 3218 * | Multipolar |  | 28 | IVF | 2 | 2 |
| 3226 (i) | NA | NO | 31 | IVF | 2 | 2 |
| 3226 (ii) * | Multipolar (dual bipolar<br>spindle) | YES | 31 | IVF | 3 | 2 |
| 3227 | Bipolar | NO | 40 | ICSI | 2 | 2 |
| 3233 (i) | Multipolar | N/A | 33 | IVF | 3 | 3 |
| 3233 (ii) * | Multipolar | YES | 33 | IVF | at least 5 | 2 |
| 3236 | Bipolar | n/a | 35 | IVF | embryo moved out of the field of imaging | embryo moved out of the field of imaging |
| 3246 * | Bipolar | NO | 33 | IVF | 2 | 2 |
| 3247 (i) - egg share * | Bipolar | NO | 29 | ICSI | 2 | 2 |
| 3247 (ii) - egg share | Bipolar | NO | 29 | ICSI | 2 | 2 |
| 3247 (iii) - egg share * | Bipolar | YES | 29 | ICSI | 2 | 2 |
| 3247 (iv) - egg share | Bipolar | YES | 29 | ICSI | 2 | 2 |

<sup>2</sup> \* denotes 16 embryos where mitosis I was filmed in its entirety

**Table S3:**
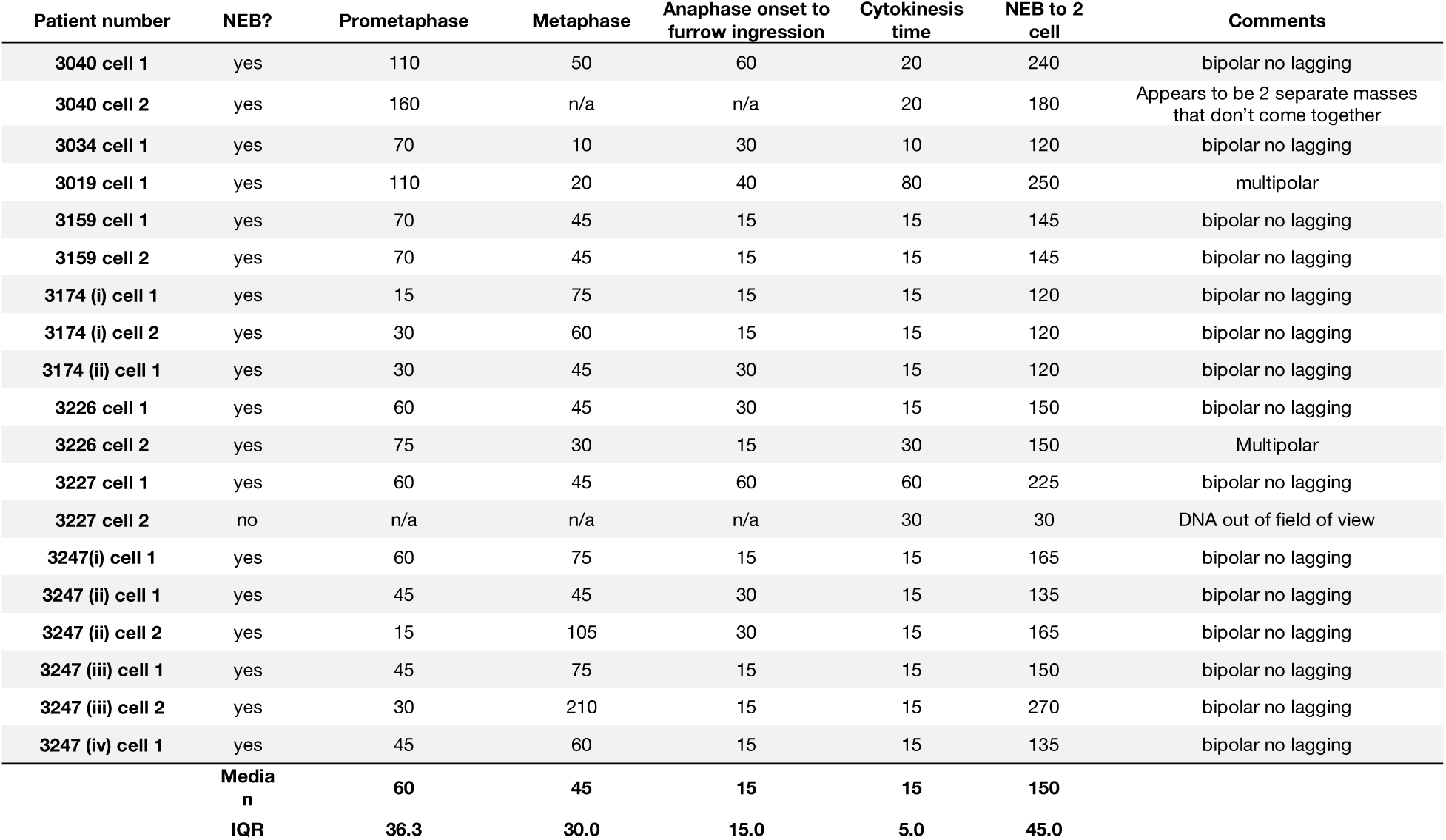
Mitosis II timing and phenotypes.

**Figure S1:**
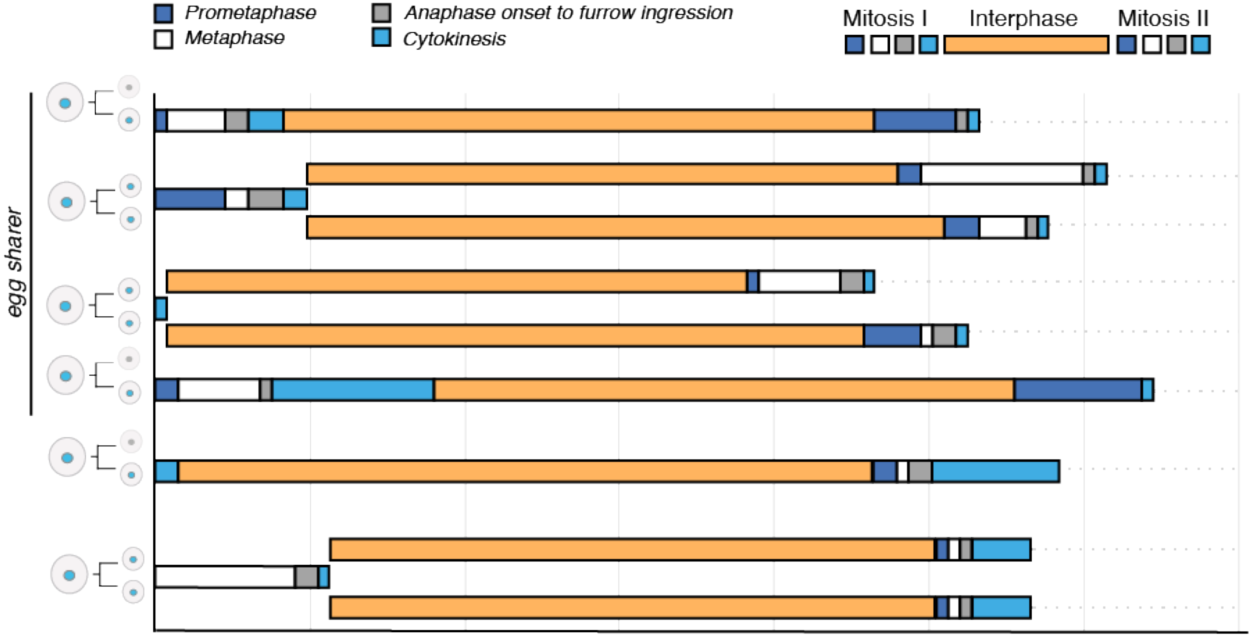
Duration of interphase between mitosis I and II of human embryos. History plots showing quantification of interphase in live-embryo movies in which both mitosis I and II were captured. Coloured bars denote timings of critical stages of mitosis I, II and interphase.

**Figure S2:**
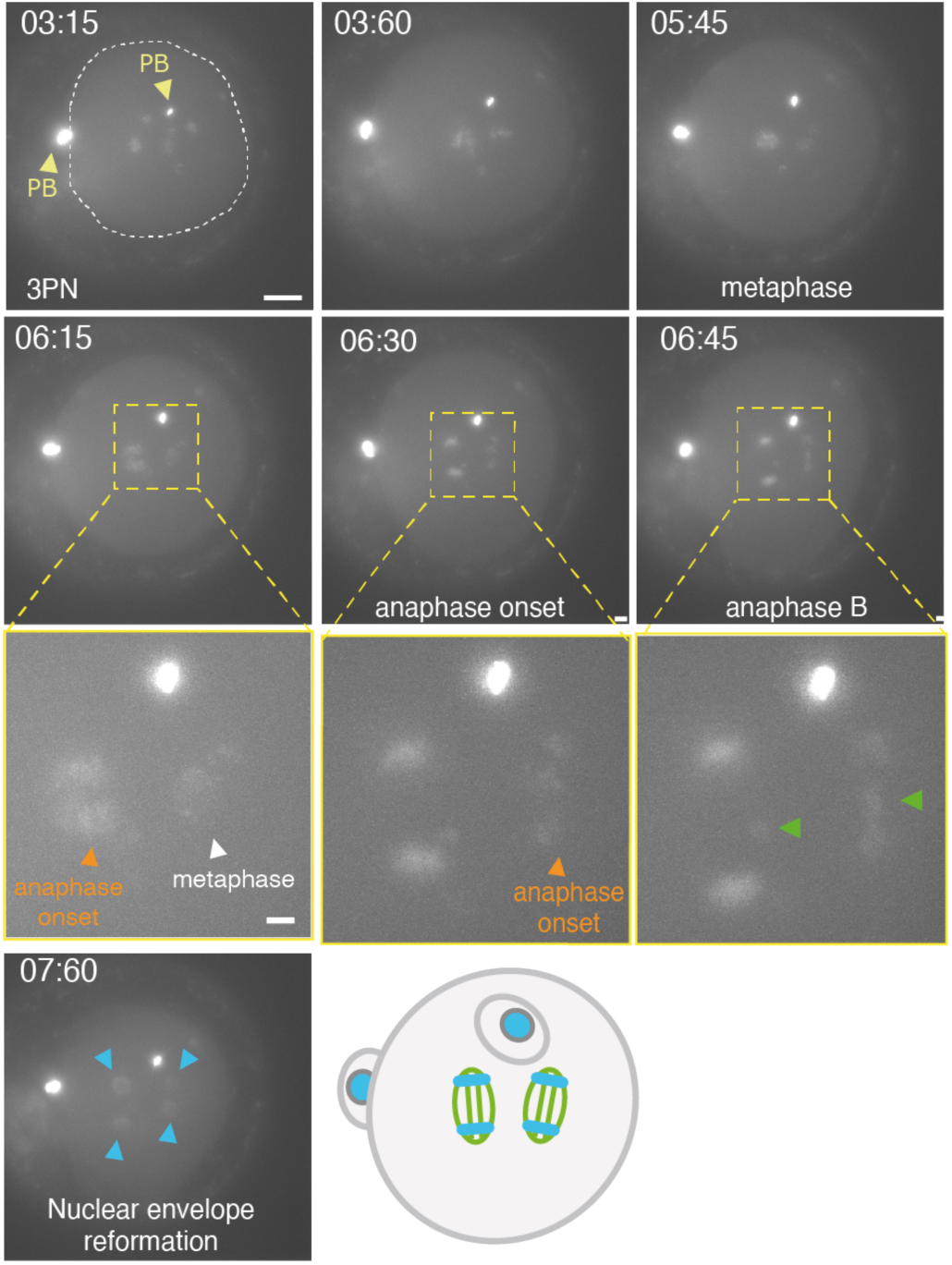
Evidence of dual spindle formation during mitosis I of a human embryo. Time lapse imaging of an IVF-fertilised 3PN human embryo progressing through mitosis I in the presence of a dual spindle. Note initiation of anaphase in left spindle before right spindle at 06:15. Green arrow heads indicate lagging chromosomes in both spindles (06:45), while blue arrow heads indicate formation of four nuclei (07:60). Chromosomes were visualised using SiR-DNA dye. Chromosome images are a maximum intensity projection of all 60 Z-slices. Time in hours:mins, scale bar: 20μm, inset 5*μ*m. *Bottom right*, schematic showing infered positon of the two spindles (and polar bodies) based on chromosome positions.

